# Genomic rearrangements uncovered by genome-wide co-evolution analysis of a major nosocomial pathogen *Enterococcus faecium*

**DOI:** 10.1101/2020.10.20.346924

**Authors:** Janetta Top, Sergio Arredondo-Alonso, Anita C. Schürch, Santeri Puranen, Maiju Pesonen, Johan Pensar, Rob J.L. Willems, Jukka Corander

**Affiliations:** Department of Medical Microbiology, University Medical Center Utrecht, Utrecht, The Netherlands; Department of Computer Science, Aalto University, FI-00076 Espoo, Finland; Department of Mathematics and Statistics, Helsinki Institute of Information Technology (HIIT), FI-00014 University of Helsinki, Finland; Pathogen Genomics, Wellcome Trust Sanger Institute, Cambridge CB10 1SA, UK; Department of Biostatistics, University of Oslo, 0317 Oslo, Norway

**Keywords:** *Enterococcus faecium*, genome-wide co-evolution analysis, genomic rearrangement

## Abstract

*Enterococcus faecium* is a gut commensal of the gastro-digestive tract, but also known as nosocomial pathogen among hospitalized patients. Population genetics based on whole-genome sequencing has revealed that *E. faecium* strains from hospitalized patients form a distinct clade, designated as clade A1 and that plasmids are major contributors to the emergence of nosocomial *E. faecium*. Here we further explored the adaptive evolution of *E. faecium* using a genome-wide co-evolution study (GWES) to identify co-evolving SNPs. We identified three genomic regions harboring large numbers of SNPs in tight linkage which are not proximal to each other based on the completely assembled chromosome of clade A1 reference hospital isolate AUS0004. Close examination of these regions revealed that they are located at the borders of four different types of large-scale genomic rearrangements, insertion sites of two different genomic islands and an IS*30*-like transposon. In non-clade A1 isolates, these regions are adjacent to each other and they lack the insertions of the genomic islands and IS*30*-like transposon. Additionally, among the clade A1 isolates there is one group of pet isolates lacking the genomic rearrangement and insertion of the genomic islands, suggesting a distinct evolutionary trajectory. *In silico* analysis of the biological functions of the genes encoded in three regions revealed a common link to a stress response. This suggests that these rearrangements may reflect adaptation to the stringent conditions in the hospital environment, such as antibiotics and detergents, to which bacteria are exposed. In conclusion, to our knowledge, this is the first study using GWES to identify genomic rearrangements, suggesting that there is considerable untapped potential to unravel hidden evolutionary signals from population genomic data.

**Impact statement:** *Enterococcus faecium* has emerged as an important nosocomial pathogen around the world. Population genetics revealed that clinical *E. faecium* strains form a distinct clade, designated as clade A1 and that plasmids are major contributors to the emergence of nosocomial *E. faecium*. Here, the adaptive evolution of *E. faecium* was further explored using an unsupervised machine learning method (SuperDCA) to identify genome-wide co-evolving SNPs. We identified three genomic regions harboring large numbers of SNPs in tight linkage which are separated by a large chromosomal distance in a clinical clade A1 reference isolate, but appeared adjacent to each other in non-clade A1 isolates. We identified four different types of large-scale genomic rearrangements and in all cases, we found insertion of two different genomic islands and an insertion element at the border. In contrast, no genomic rearrangement and insertions were identified among a group of clade A1 pet isolates, suggesting a distinct evolutionary trajectory. Based on the *in silico* predicted biological functions, we found a common link to a stress response for the genes encoded in three regions. This suggests that these rearrangements may reflect adaptation to the stringent conditions in the hospital environment, such as antibiotics and detergents, to which bacteria are exposed.

**Data summary:** Raw core-genome alignment (1.1 MB, Harvest suite v1.1.2), including the 1,644 Clade A isolates and the complete *E. faecium* AUS0004 (accession number CP003351) as a reference is available under the following gitlab repository https://gitlab.com/sirarredondo/efm_gwes.

## Introduction

*Enterococcus faecium* are commensals of the gastrointestinal tract but are now recognized as a major causative agent of healthcare associated infections (1). The transition from commensal to nosocomial pathogen coincided with increased resistance to antibiotics (2). Since the early 1980s, *E. faecium* first gained high-level resistance to ampicillin, followed by resistance to aminoglycosides, fluoroquinolones and glycopeptides, particularly vancomycin (3,4). Previous whole genome sequencing (WGS) studies identified a split in the *E. faecium* population into two lineages, including a hospital-associated clade (clade A) and a community-related clade (clade B) (5,6). Later, clade A was subdivided into clade A1 representing the majority of hospital associated isolates and clade A2 for animal-related isolates (7), although two studies which included a larger collection of animal-related isolates suggested that these isolates clustered in polyphyletic groups and not in a distinct clade A (8,9). In a recent study also including 55 isolates from Latin America, phylogenomics suggests that the animal isolates represent multiple lineages that diverged prior to the emergence of the clinical subclades in clade A (10).

Recently, we determined the plasmid content (plasmidome) of 1,644 *E. faecium* isolates from different sources, countries and years using short- and long-read whole genome sequencing technologies in combination with machine-learning classifiers (9,11). This analysis revealed that the hospital-associated isolates carried a larger number of plasmid sequences compared to isolates from other sources and different configurations of plasmidome populations in the hospital environment. In addition, the source specificity was determined for the chromosomal, plasmid and whole-genome components. It was concluded that plasmid sequences have the highest contribution to source specificity.

In this paper, WGS of the same 1,644 isolates were used with the aim to identify signals of selection acting to shape co-evolution of SNPs. Co-evolved SNPs can facilitate adaptation to different environments when sequentially selected mutations in adaptive elements decrease the actual fitness costs of individual mutations (antagonistic epistasis) and thus have a beneficial effect on fitness (12). For this purpose, we used the Direct Coupling Analysis tool (SuperDCA) as previously described for *Streptococcus pneumoniae* where previously undetected epistatic interactions related to e.g. survival of the pneumococcus at lower temperatures were discovered (13). For this study, SuperDCA was applied on the core genome alignment of *E. faecium* to identify likely candidates of sequentially selected or coupled mutations not in strong linkage disequilibrium due to chromosomal proximity(13).

## Results

### Co-evolution of core genomic variation

SuperDCA was applied to a core-genome alignment generated using the Harvest suite, including 1,644 Clade A isolates (S1 Table, accession number PRJEB28495)(9) and the complete *E. faecium* genome of strain AUS0004 (accession number CP003351) as reference (14). All locus tags mentioned below refer to AUS0004. SuperDCA identified 262,877 significant couplings between SNPs with a distance of > 1 kbp (S2 Table). We generated a top-10 list of SNP positions with the highest number of links using non-overlapping bins of 100 bp (S1A and B Fig). To obtain insight into the distribution on the AUS0004 core genome, all links for each of these ten SNP positions was plotted (S1C Fig). In this paper, we will mainly focus on the SNP positions from bins 3 and 10 (S1B Fig) representing genes *EFAU004_02176* and *EFAU004_02173*, because we were able to identify a possible/plausible biological explanation. For both bins, we observed a similar pattern of coupled SNPs with the genes *EFAU004_00665* to *EFAU004_00675* located at a large chromosomal distance of around 500 kbp in the AUS0004 core genome (S1C Fig). Detailed examination of the total list of coupled SNPs revealed similar patterns of coupled SNPs for a larger region encompassing genes *EFAU004_02173* to *EFAU004_02178*, referred to region-1 (Fig 1A). There were in total 2,323 coupled SNP positions, i.e. between 181 different SNP positions from region-1 and 131 different SNP positions in genes *EFAU004_00665* to *EFAU004_00675*, referred to region-2 (Fig 1A and S1C and/or S3 Table). In addition, all coupled SNPs for region-1 and -2 were retrieved from the total list. These 17,236 linked SNPs were plotted on the core genome to obtain insight into the distribution of the links (Fig 1A). This revealed a cluster of in total 142 linked SNPs, represented by 32 different SNP positions in region-2 and 25 SNP positions in genes *EFAU004_02092* to *EFAU004_02100*, representing region-3, located close to region-1 (Fig 1A and S3 and/or S4 Table).

**Fig 1.**
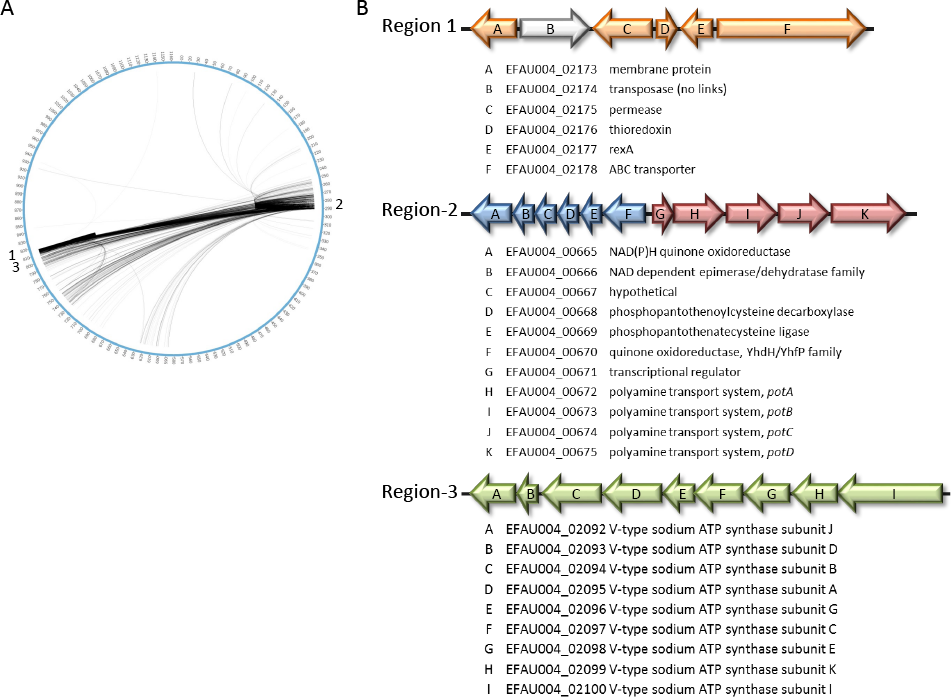
A: All 17,236 epistatic links of the genes contained in region-1, -2 and -3. B: genomic organization and annotation of genes from region 1-3 in *E. faecium* AUS0004.

In order to elucidate a possible biological explanation for the identified links between region-1, -2 and -3, we first determined putative biological functions for the proteins in these regions using homology search for similarity with other proteins and investigate domains with known function.

### Putative biological functions for proteins in region-1

Region-1 contains five protein encoding genes and a transposase with no links (Fig 1B). For two proteins a function prediction is challenging, i.e. EFAU004_02173 is annotated as a membrane protein, but there are only domains of unknown function (DUF), while EFAU004_02175 is annotated as a permease with unknown specificity. In contrast, EFAU004_02176 contained conserved domains belonging to the thioredoxin-like family (Trx-like) and EFAU004_02177 contained conserved domains belonging to the Rex (Rex-like) family of transcriptional regulators (Fig 1B). Trx-like and Rex-like are likely involved in redox homeostasis of the bacterial cell, which was found to be critical for DNA synthesis and defense against oxidative stress (15). The exact function of the ABC transporter (EFAU004_02178) is difficult to predict despite the presence of several domains. The protein contains two so-called AAA domains and a leucine-zipper (bZIP) domain. Proteins with AAA domains are members of a conserved family of ATP-hydrolyzing proteins with all kind of activities in many cellular pathways, including replication, DNA and protein transport, transcriptional regulation, ribosome biogenesis, membrane fusion, and protein disaggregation or degradation (16). bZIP domains are known to be involved in transcriptional regulation, e.g. in stress conditions like heat in *Salmonella* (17,18), or abiotic stress in eukaryotes, e.g. plants (19). Furthermore, homology search against the GenBank database to non-*Enterococcaceae* revealed co-localization of a similar ABC transporter and *rexA* gene in other species like *Streptococcus pneumoniae* (strain 2842STDY5753625, accession number FEGY01000003), *Streptococcus agalactiae* (strain DK-PW-092, accession number LBKE01000082) and *Listeria monocytogenes* (strain CFSAN060067, accession number AABAZK010000087), suggesting that this ABC transporter might also be involved in some kind of stress response.

### Putative biological functions for proteins in region-2

The genes from region-2 can be split into two distinct gene clusters based on their putative biological functions (Fig 1B). The first cluster encompasses genes *EFAU004_00665* to *EFAU004_00670* which, based on their annotation, are putatively involved in respiration and coenzyme A biosynthesis (Fig 1B). Although SNPs linked with region-1 were observed with all these genes, we will focus on two specific genes, i.e. *EFAU004_00665* and *EFAU004_00670* as they contained the majority of linked SNPs (S3 and S4 Table). Very similar conserved domains for NAD(P)H:quinone oxidoreductase were identified in both proteins. EFAU004_00670 contains transmembrane helices and is therefore likely membrane bound, while EFAU004_00665 lacks these transmembrane helices and is therefore likely to be soluble. The conserved domains in EFAU004_00670 had a high similarity to YhdH of *Escherichia coli* (overall amino acid similarity of 48%) The conserved domains in EFAU004_00665 was very similar to QorEc of *E. coli* (overall amino acid similarity of 32%) and QorTt *Thermus thermophiles* (overall amino acid similarity of 32%). In *Staphylococcus aureus*, expression of a gene cluster containing two *qor*-like genes was induced under oxidative stress conditions (20). These two Qor proteins (SA1988 and SA1989) are predicted as soluble and SA1988 contained similar conserved domains with an overall amino acid similarity of 26% with EFAU004_00665. This could suggest that this protein could be involved in stress response as well.

The second gene cluster contains genes *EFAU004_00671* to *EFAU004_00675* and is likely organized as an operon (Fig 1B). EFAU004_00671 contains a helix-turn-helix motif and is therefore predicted as transcriptional regulator. Its location upstream of *EFAU004_00672* to *EFAU004_00675* suggests that it regulates the expression of these genes. EFAU004_00672 to EFAU004_00675 display similarity with a polyamine transport system as described for *Streptococcus pneumoniae*, including an ATP binding protein PotA, 2 permeases PotB and PotC and a substrate binding protein PotD (21). Polyamines are polycationic molecules and are required for optimal growth in both eukaryotic and prokaryotic cells and are implicated in pathogenicity of *S. pneumoniae*. In *S. pneumoniae* polyamines are pivotal in survival strategies in the host when bacteria are confronted with stress conditions like temperature shock, oxidative stress, or choline limitation (21).

### Putative biological functions for proteins in region-3

Genes *EFAU004_02092* to *EFAU004_02100* are annotated to encode a membrane bound V-type ATPase (Fig 1B). An identical V-type ATPase has been studied in detail in *Enterococcus hirae* (100% amino acid identity) (22). The V-type ATPase belongs to the family of proton pumps and are involved in the translocation of Na+ or H+ over the cell membrane by using the energy of ATP. In *E. hirae* the V-type ATPase appeared highly expressed under stress conditions like high pH and plays an important role in sodium homeostasis under these conditions.

### Genomic rearrangements in completed *E. faecium* genomes

For 48 of 1,644 isolates, a fully assembled circular chromosome was available (accession number PRJEB28495) as described previously (11). This allowed us to assess the genomic organization in these isolates. Using the non-clade A1 isolate E0139 as the reference, four different types of large chromosomal rearrangements were observed among 38 isolates. Type-1 (n = 1) involves an inversion of 0.73 Mbp; type-2 (n = 34) an inversion of 1.2 Mbp; type-3 (n = 2) two inversions of 0.38 Mbp; and type-4 (n = 1) an inversion of 0.12 Mbp (Fig 2A-D and S2 Fig). Except for the type-1 inversion, all these genomic rearrangements were exclusively observed in clade A1 isolates (Fig 3). In clade A1 isolates we detected two previously described genomic islands, inserted adjacent to the genomic rearrangements that were described as being enriched among clade A1 hospital isolates (Fig 2B-D) (23,24). One genomic island, putatively encoding a carbohydrate transport and metabolism pathway (23), was always found to be located downstream of the ABC transporter (*EFAU004_02178*) of region-1 (Fig 4A). The other island encoding a phosphotransferase system (PTS) (24) was always found to be located downstream *potD* (*EFAU004_00675*) of the polyamine transport system of region-2 (Fig 4A). In all cases, the genomic rearrangement was flanked by an IS*30*-like transposon downstream of a methionine synthase (*metE*) and a pyridine nucleotide-disulfide oxidoreductase (*pyr*) (Fig 4A). In the 10 isolates for which a fully assembled circular chromosome was available and that lacked the genomic rearrangement, insertions of both genomic islands and IS*30*-like transposon, region-1 and region-2 are located adjacent to each other as schematically indicated for the non-clade A1 isolate *E. faecium* E0139 (Fig 4B).

**Fig 2.**
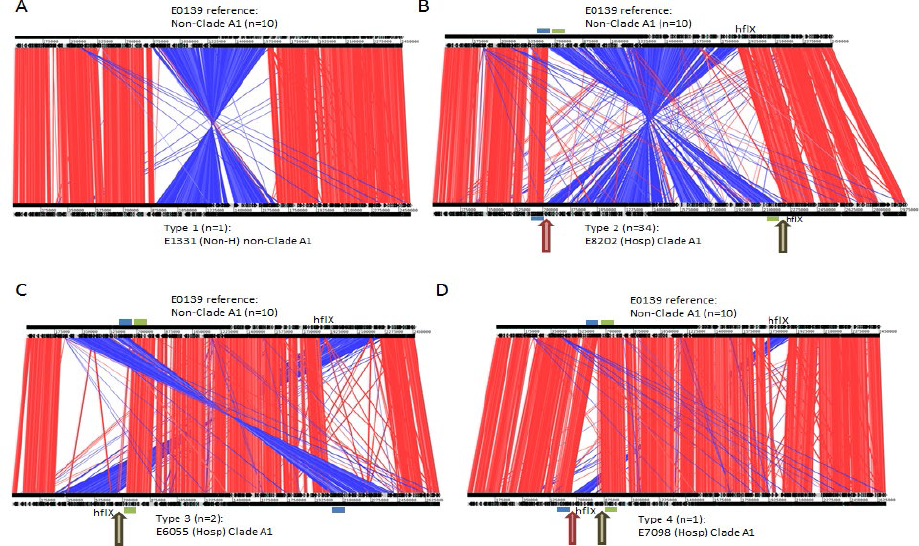
Genome comparisons using strain E0139 (non-clade A1) as a reference. A: E0139 compared to type 1 strain E1334; B, E0139 compared to type 2 strain E8202; C, E0139 compared to type 3 strain E6055; D, E0139 compared to type 4 strain E7098. “n” indicates the number of isolates with a similar genomic organization and for which a complete chromosome was available Same-strand DNA similarity is shaded red, while reverse similarity is shaded blue. Blue bar: indication position *pot* operon; Green bar: indication position ABC transporter; red arrow: insertion site phosphotransferase system (PTS) encoding genomic island [24]; brown arrow: insertion site carbohydrate transport system encoding genomic island [23].

**Fig 3.**
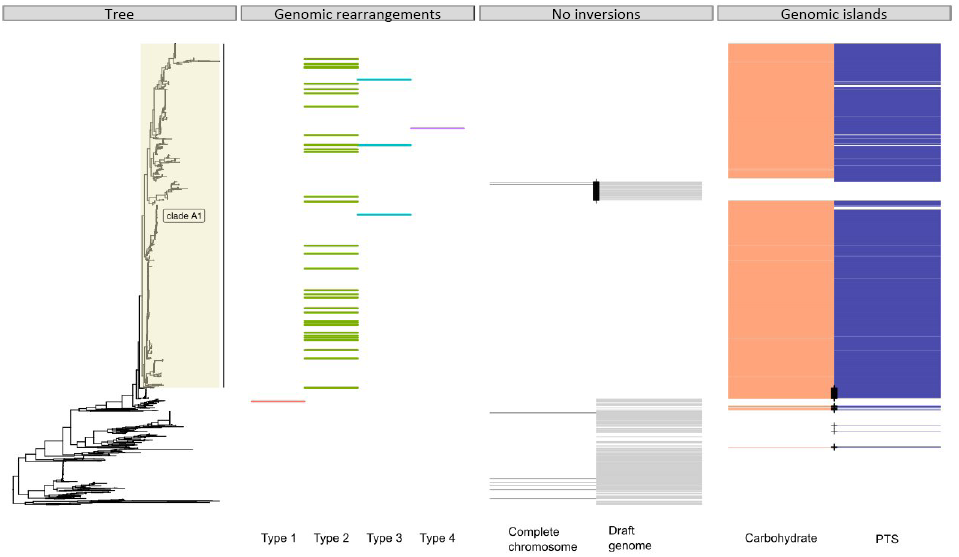
Core genome tree based on 1644 strains with the three metadata panels: i) distribution of genomic rearrangements types among 38 complete chromosomes, ii) indication of strains lacking a genomic rearrangement among 10 complete chromosomes and draft genomes and iii) indication of strains with insertion of a carbohydrate transport system encoding genomic island, orange [23] or a phosphotransferase system (PTS) encoding genomic island, purple[24]. Black cross: indication of clade A1 dog isolates that lack the insertions/inversion or non-clade A1 dog isolates that do contain insertions/inversion.

**Fig 4.**
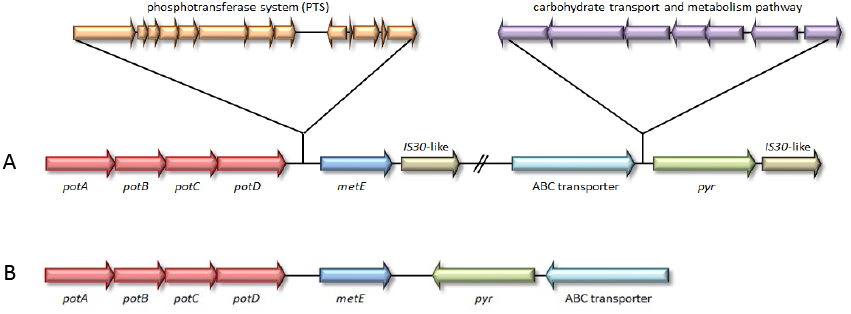
A: Genomic organization of the most predominant type 2 genomic rearrangement in strain E8202 with indication of insertion and inversion site. B: Genomic organization for reference strain E0139.

We determined the presence of the two genomic islands and their insertion sites in the draft genomes of the remaining 1,596 isolates and confirmed that the genomic islands were always inserted at the same position in 93% of the clade A1 isolates (Fig 3). In addition, the insertions were also identified in 57 (14%) non-clade A1 isolates, which mainly represented dog isolates (n = 43) (Fig 3). In contrast, a branch in clade A1 consisting of 68 genetically closely related isolates, including 55 pet, 9 isolates from hospitalized patients and 4 from non-hospitalized persons, lacked both genomic island insertions (Fig 3). Based on these draft genomes, we cannot determine whether or which genomic rearrangement could have occurred, though we always observed a contig break downstream of *metE* and *pyr* strongly suggesting the presence of an IS element on that position. In isolates that lacked insertions of the two genomic islands the genomes were organized in a manner comparable to the non-clade A1 isolate E0139, i.e. without inversion (Fig 3 and 4B).

## Discussion

In this study, we identified large genomic rearrangements, enriched in *E. faecium* hospital-associated clade A1 strains, which were uncovered by the appearance of co-evolved SNPs in three chromosomal regions; region-1, region-2 and region-3. These chromosomal regions were located on the borders of the genomic rearrangement and therefore adjacent to each other in the majority of non-clade A1 isolates but at a large distance in clade A1 isolates. In all cases, the genomic rearrangement is flanked by an IS*30*-like transposon downstream *metE* and *pyr*, which suggests that the rearrangement is the result of recombination between two IS*30*-like elements. In addition, upstream of *metE* and *pyr*, we identified the insertion of two, clade A1 enriched, genomic islands, encoding a PTS and carbohydrate transport system, respectively (23,24). For both genomic islands, it has been suggested that they may provide a fitness advantage, which was confirmed for PTS in an *in vivo* mouse colonization model (24). Recently, Yan *et al*. reconstructed the evolution of the marine photosynthetic microorganism *Prochlorococcus* based on genome rearrangements and identified rearrangement hotspots which were all in the vicinity of genomic islands (25). In fact, the authors suggest that the genomic islands serve as hotspots that induce genome rearrangement. In addition, they observed that different clades shared a conserved backbone, but also contained clade specific regions, which were associated with ecological adaptations. Also for *Pseudomonas putida*, a gram-negative bacterium that can be found in different environments, it was shown that horizontal gene transfer played a key role in adaptation process, as many of the niche-specific functions were found to be encoded on clearly defined genomic islands and were linked to genomic rearrangements (26). The observation that among the clade A1 isolates only the pet isolates do not contain the two genomic islands and genomic rearrangement, may suggest that these elements are not advantageous in the niche represented by the gut of pet animals. The predominant chromosomal rearrangement, type-2, identified in 71% of the fully assembled genomes from the current study is also present in other publicly available completed genomes such as *E. faecium* DO (NC_017960.1) (27), *E. faecium* Aus0085 (NC_021994.1) (28) or *E. faecium* E39 (NZ_CP011281.1). For the isolates in our study for which we only have a draft genome, it is difficult to determine whether genomic rearrangements have occurred. However, in isolates in which *metE* or *pyr* were not adjacent to each other as in the non-clade A1 configuration, we always identified an IS*30*-like element downstream the *metE* or *pyr* gene at the border of a contig suggesting a potential genomic rearrangement. A chromosomal rearrangement in *E. faecium* was also observed in the first completely sequenced isolate AUS0004, where it was hypothesized that it occurred between 2 phage elements resulting in a replichore imbalance (14). However, we did not observe a replichore imbalance in any of the 38 completely assembled genomes with large chromosomal rearrangements (data not shown), which suggests that AUS0004 is an exception in that regard.

Genomic rearrangements or inversions are not rare among prokaryotic genomes as was investigated by Repar *et al*. (29). The authors developed an alignment-based approach to systematically assess inversion symmetry between pairs of prokaryotic genomes, including the genera *Bacillus* and *Clostridium*, which belong to the phylum Firmicutes like *E. faecium*. *Bacillus* and *Clostridium* are dominated by X-shaped inversions that are symmetric around the *ori-ter* axis similar to the type 1-3 inversions from this study.

The gene clusters located in the chromosomal regions-1, -2, and -3 that harbored co-evolved SNPs clearly encode common functions. The thioredoxin pathway, Rex-like (region-1) and the NAD(P)H dependent oxidoreductase/epimerase proteins (region-2) are all dependent on the oxidation/reduction of NAD+/NADH and involved in redox homeostasis, while the polyamine transport system (region-2) and V-type ATPase (region-3) are both involved in ATP hydrolysis/synthesis. The fact that in non-clade A1 isolates chromosomal regions-1, -2 and -3 are located adjacent to each other explains the high number of statistically significantly linked SNPs between the genes in these two regions, as they likely have co-evolved in fairly tight linkage disequilibrium (LD). Conversely, this finding illustrates that the co-evolutionary analysis of SNPs can uncover inversions in the chromosome. Enterococci are rich in mobile genetic elements harbouring IS elements and transposons, many of which can integrate into the chromosome (REF) and may drive the observed rearrangements. These rearrangements may play a role in adaptation to new environments, more particularly adaptation to stress conditions, like the hospital environment where bacteria are exposed to different stressors including high concentrations of antibiotics, detergents and antiseptics. Furthermore, it is possible that the rearrangements are associated with the rapid adaptation of the organism to survive in the perturbed microbiota of hospitalized patients and that survival in the overall context of the intestinal microbiome may be an important driver of evolution. The lack of comparable selection pressure is a plausible explanation of the observed absence of genomic arrangements among the pet isolates from clade A1.

To our knowledge, identification of genomic rearrangements by direct coupling analysis of SNPs has not been previously described in the literature. The expected rapid increase in the availability of long-read sequences combined with large population-based collections of short-read sequenced genomes that are amenable to a statistical LD analysis such as done here will thus facilitate the development of new methods to identify candidates of inversions associated with selective advantage.

## Material and Methods

### Genomic DNA Sequencing and assembly

A detailed description of Illumina and ONT sequencing has been described previously and includes a full description on ONT selection of *E. faecium* isolates (n = 62) (9,11) and consecutive hybrid assembly using Unicycler (30). The R package ggtree (version 1.14.6) (31) was used to plot distinct metadata panels together with a previously described core genome tree of 1,644 *E. faecium* strains (9).

### Co-evolution analysis

A core-genome alignment (1.1 MB) was generated using the Harvest suite v1.1.2, including the 1,644 Clade A isolates and the complete *E. faecium* AUS0004 (accession number CP003351 (14)) as reference. SuperDCA was performed with the default setting as described by Puranen *et al*. (13) using the alignment of 1,644 isolates which was unfiltered for recombination events. Circos was used for the visualization of coupled SNPs on the genome (32).

### Prediction of putative biological functions of proteins from region 1-3

To predict putative biological functions of proteins, the protein sequences were compared with the non-redundant public database using the National Center for Biotechnology Information (NCBI) BLASTp server (www.ncbi.nlm.nih.gov/BLAST/) (33). In addition, the graphic summary was used to determine whether putative conserved domains were identified and if present this graphical summary was selected to investigate in more detail the list of conserved domains in order to make a prediction of the putative biological function of the protein.

### *E. faecium* genome organization

To unravel rearrangements in the chromosome, we considered a non-clade A1 isolate from a non-hospitalized person (E0139) with a complete chromosome sequence as reference. We only considered isolates with a complete circular chromosome (n = 48). Pairwise chromosomal comparisons were computed using blastn (version 2.7.1+) and alignments were visualized using the Artemis Comparison tool (version 17.0.1). The insertions of two genomic islands encoding a PTS (24) and carbohydrate transport system (23) and their putative co-localization with the *pot* operon and ABC transporter, respectively was determined by BLAST. Additionally, for each chromosomal rearrangement, we generated dotplots using Gepard (version 1.40) (34) using the non-clade A1 isolate E0139 as a reference. Clone manager 9 was used to visualize the genomic organization of region 1-3.

## Author statements

### Authors and contributors

Conceptualization, JC and RJLW; software, SP, MP and JP; formal analysis SA, JT and ACS; writing of original draft, JT, SA, ACS, JC and RJLW; funding acquisition, RJLW and JC.

## Supporting information

**S1 Fig.**
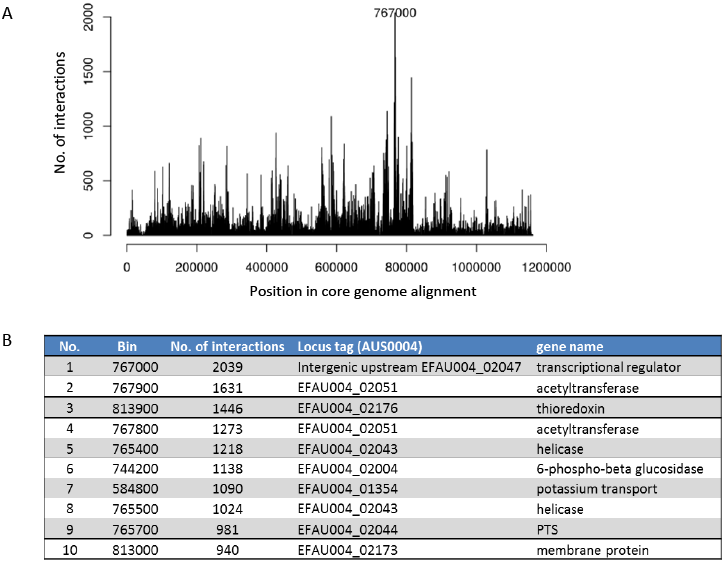
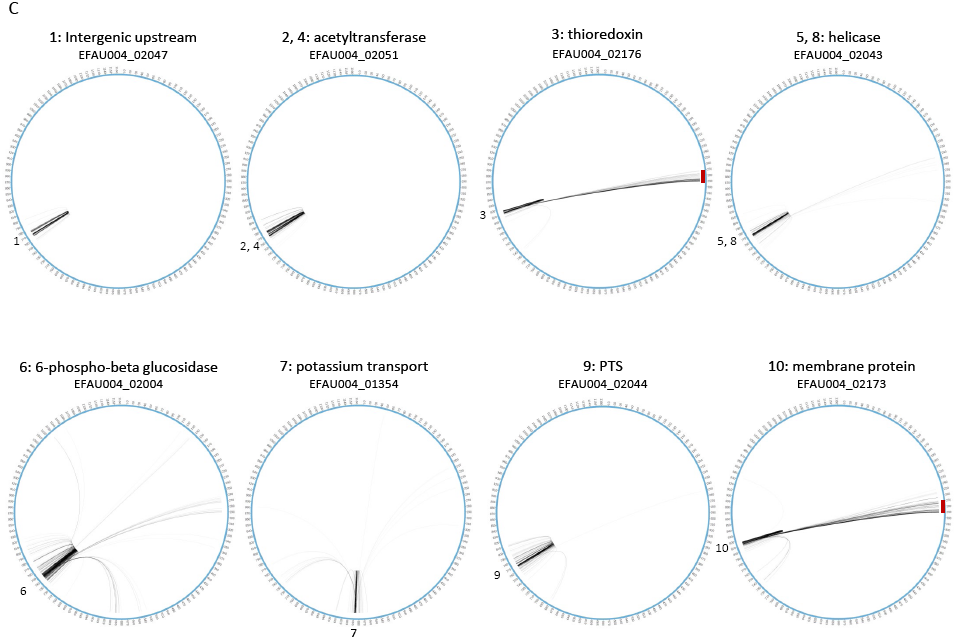
Selection procedure for the highest number of coupled SNP positions. A: histogram depicting the number of links per non-overlapping bin of 100-bps in the core genome alignment. B: Top-10 list of the 100-bps bins with the highest number of links. C: Circos plots depicting all coupled SNP positions for the Top-10 100-bps bins. Numbers refer to the ranking position. Gene names and locus tags refer to the *E. faecium* AUS0004 reference genome. The red bar for thioredoxin and the membrane protein indicate the overlap in interactions.

**S2 Fig.**
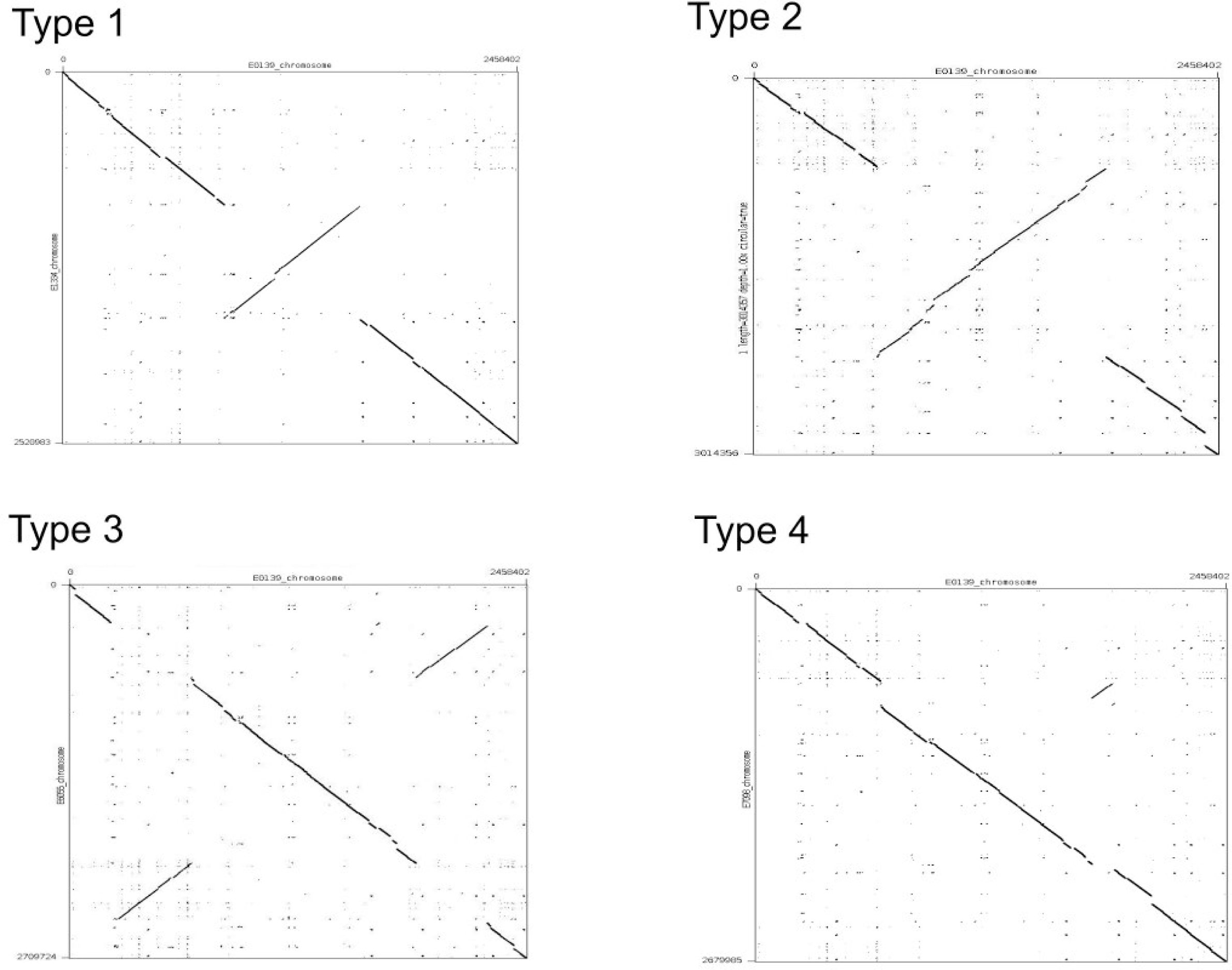
Dotplots of four different types of genomic rearrangements observed in our set of 48 complete chromosome sequences using strain E0139 (non-clade A1) as reference. Comparison with strain: E1334 for type 1, E8202 for type 2, E6055 for type 3 and E7098 for type 4.

**S1 Table. Metadata 1,644 clade A strains**

**S2 Table. List of significant couplings between SNPs with a distance of > 1 kbp**

**S3 Table. List of coupled SNP positions between region-1 and region-2**

**S4 Table. List of coupled SNP positions between region-2 and region-3**

## References

1. Weiner LM, Webb AK, Limbago B, Dudeck MA, Patel J, Kallen AJ, et al. Antimicrobial-Resistant Pathogens Associated with Healthcare-Associated Infections: Summary of Data Reported to the National Healthcare Safety Network at the Centers for Disease Control and Prevention, 2011-2014. Infect Control Hosp Epidemiol. 2016;37(11):1288–301.

2. Lebreton, F., Willems, R.J.L. and Gilmore MS. *Enterococcus* Diversity, Origins in Nature, and Gut Colonization. In: Enterococci: From Commensals to Leading Causes of Drug Resistant Infection [Internet]. 2014. p. 1–56. Available from: http://www.ncbi.nlm.nih.gov/pubmed/24649513

3. Gilmore MS, Lebreton F, van Schaik W. Genomic transition of enterococci from gut commensals to leading causes of multidrug-resistant hospital infection in the antibiotic era. Vol. 16, Current Opinion in Microbiology. 2013. p. 10–6.

4. Guzman Prieto AM, van Schaik W, Rogers MRC, Coque TM, Baquero F, Corander J, et al. Global emergence and dissemination of enterococci as nosocomial pathogens: Attack of the clones? Vol. 7, Frontiers in Microbiology. 2016.

5. Galloway-Peña J, Roh JH, Latorre M, Qin X, Murray BE. Genomic and SNP analyses demonstrate a distant separation of the hospital and community-associated clades of *Enterococcus faecium*. PLoS One. 2012;7(1).

6. Palmer KL, Godfrey P, Griggs A, Kos VN, Zucker J, Desjardins C, et al. Comparative genomics of enterococci: Variation in *Enterococcus faecalis*, clade structure in *E. faecium*, and defining characteristics of *E. gallinarum* and *E. casseliflavus*. MBio. 2012;3(1):1–11.

7. Lebreton F, van Schaik W, McGuire AM, Godfrey P, Griggs A, Mazumdar V, et al. Emergence of epidemic multidrug-resistant *Enterococcus faecium* from animal and commensal strains. MBio. 2013;4(4).

8. Raven KE, Reuter S, Reynolds R, Brodrick HJ, Russell JE, Török ME, et al. A decade of genomic history for healthcare-associated *Enterococcus faecium* in the United Kingdom and Ireland. Genome Res. 2016;26(10):1388–96.

9. Arredondo-Alonso S, Top J, McNally A, Puranen S, Pesonen M, Pensar J, et al. Plasmids shaped the recent emergence of the major nosocomial pathogen *Enterococcus faecium*. MBio. 2020;11(1):1–17.

10. Rios R, Reyes J, Carvajal LP, Rincon S, Panesso D, Echeverri AM, et al. Genomic Epidemiology of Vancomycin-Resistant *Enterococcus faecium* (VREfm) in Latin America: Revisiting The Global VRE Population Structure. Sci Rep [Internet]. 2020;10(1):5636. Available from: http://dx.doi.org/10.1038/s41598-020-62371-7

11. Arredondo-Alonso S, Rogers MRC, Braat JC, Verschuuren TD, Top J, Corander J, et al. Mlplasmids: a User-Friendly Tool To Predict Plasmid- and Chromosome-Derived Sequences for Single Species. Microb genomics [Internet]. 2018;4(11). Available from: http://www.microbiologyresearch.org/content/journal/mgen/10.1099/mgen.0.000224.v1

12. Lagator M, Colegrave N, Neve P. Selection history and epistatic interactions impact dynamics of adaptation to novel environmental stresses. Proc R Soc B Biol Sci [Internet]. 2014;281. Available from: https://pubmed.ncbi.nlm.nih.gov/25232137

13. Puranen S, Pesonen M, Pensar J, Xu YY, Lees JA, Bentley SD, et al. SuperDCA for genome-wide epistasis analysis. Microb genomics. 2018;4(6).

14. Lam MMC, Seemann T, Bulach DM, Gladman SL, Chen H, Haring V, et al. Comparative analysis of the first complete *Enterococcus faecium* genome. J Bacteriol. 2012;194(9):2334–41.

15. Wang E, Bauer MC, Rogstam A, Linse S, Logan DT, Von Wachenfeldt C. Structure and functional properties of the *Bacillus subtilis* transcriptional repressor Rex. Mol Microbiol. 2008;69(2):466–78.

16. Elsholz AKW, Birk MS, Charpentier E, Turgay K. Functional diversity of AAA+ protease complexes in *Bacillus subtilis*. Front Mol Biosci. 2017;4(JUL):1–15.

17. Hurme R, Berndt KD, Namork E, Rhen M. DNA binding exerted by a bacterial gene regulator with an extensive coiled-coil domain. J Biol Chem. 1996;271(21):12626–31.

18. Hurme R, Berndt KD, Normark SJ, Rhen M. A proteinaceous gene regulatory thermometer in *Salmonella*. Cell. 1997;90(1):55–64.

19. Alves MS, Dadalto SP, Gonçalves AB, de Souza GB, Barros VA, Fietto LG. Transcription factor functional protein-protein interactions in plant defense responses. Proteomes. 2014;2(1):85–106.

20. Maruyama A, Kumagai Y, Morikawa K, Taguchi K, Hayashi H, Ohta T. Oxidative-stress-inducible *qorA* encodes an NADPH-dependent quinone oxidoreductase catalysing a one-electron reduction in *Staphylococcus aureus*. Microbiology. 2003;149(2):389–98.

21. Shah P, Romero DG, Swiatlo E. Role of polyamine transport in *Streptococcus pneumoniae* response to physiological stress and murine septicemia. Microb Pathog. 2008;45(3):167–72.

22. Murata T, Kawano M, Igarashi K, Yamato I, Kakinuma Y. Catalytic properties of Na+-translocating V-ATPase in *Enterococcus hirae*. Biochim Biophys Acta - Bioenerg. 2001;1505(1):75–81.

23. Heikens E, Van Schaik W, Leavis HL, Bonten MJM, Willems RJL. Identification of a novel genomic island specific to hospital-acquired clonal complex 17 *Enterococcus faecium* isolates. Appl Environ Microbiol. 2008;74(22):7094–7.

24. Zhang X, Top J, De Been M, Bierschenk D, Rogers M, Leendertse M, et al. Identification of a genetic determinant in clinical *Enterococcus faecium* strains that contributes to intestinal colonization during antibiotic treatment. J Infect Dis. 2013;207(11):1780–6.

25. Yan W, Wei S, Wang Q, Xiao X, Zeng Q, Jiao N, et al. Genome rearrangement shapes *Prochlorococcus* ecological adaptation. Appl Environ Microbiol. 2018;84(17):e01178–18.

26. Wu X, Monchy S, Taghavi S, Zhu W, Ramos J, van der Lelie D. Comparative genomics and functional analysis of niche-specific adaptation in *Pseudomonas putida*. FEMS Microbiol Rev. 2011;35(2):299–323.

27. Qin X, Galloway-P?a JR, Sillanpaa J, Roh JH, Nallapareddy SR, Chowdhury S, et al. Complete genome sequence of *Enterococcus faecium* strain TX16 and comparative genomic analysis of *Enterococcus faecium* genomes. BMC Microbiol [Internet]. 2012;12(1):1. Available from: BMC Microbiology

28. Lam MMC, Seemann T, Tobias NJ, Chen H, Haring V, Moore RJ, et al. Comparative analysis of the complete genome of an epidemic hospital sequence type 203 clone of vancomycin-resistant *Enterococcus faecium*. BMC Genomics [Internet]. 2013;14(1):1. Available from: BMC Genomics

29. Repar J, Warnecke T. Non-random inversion landscapes in prokaryotic genomes are shaped by heterogeneous selection pressures. Mol Biol Evol. 2017;34(8):1902–11.

30. Wick RR, Judd LM, Gorrie CL, Holt KE. Unicycler: Resolving bacterial genome assemblies from short and long sequencing reads. PLoS Comput Biol. 2017;13(6):1–22.

31. Yu G, Smith DK, Zhu H, Guan Y, Lam TTY. GGTREE: an r package for visualization and annotation of phylogenetic trees with their covariates and other associated data. Methods Ecol Evol. 2017;8:28–36.

32. Krzywinski M, Schein J, Birol I, Connors J, Gascoyne R, Horsman D, et al. Circos: An information aesthetic for comparative genomics. Genome Res. 2009;19(9):1639–45.

33. Altschul S. Basic Local Alignment Search Tool. J Mol Biol. 1990;215(3):403–10.

34. Krumsiek J, Arnold R, Rattei T. Gepard: A rapid and sensitive tool for creating dotplots on genome scale. Bioinformatics. 2007;23(8):1026–8.

